# Adolescent gulls have the opportunity for social development at breeding colonies

**DOI:** 10.1101/2023.05.26.542504

**Authors:** Liam U. Taylor

## Abstract

Most seabirds delay reproduction for multiple years. The long-standing ecological hypothesis is that seabirds delay reproduction until they pass a foraging efficiency threshold. This foraging development hypothesis is puzzling for seabirds with progressive delayed plumage maturation, such as American Herring Gulls (Larus argentatus smithsonianus). Young American Herring Gulls pass through a distinct series of predefinitive plumages in their early years, suggesting a process of adolescence rather than a binary switch between energetic immaturity and maturity. Drawing on facts from both colonial seabirds and tropical lekking birds, I propose an additional life history hypothesis: young gulls undergo a phase of social development—rather than foraging development alone—during which time predefinitive plumages function as signals that reduce the costs of social engagement at breeding colonies. I tested one facet of the hypothesis: predefinitive gulls have an opportunity for social development at colonies before breeding. A unique prediction is that predefinitive gulls are common at breeding colonies, socially engaged, and not breeding. I conducted counts, quantitative tests of territoriality and conflict, and qualitative behavioral observations of American Herring Gulls at two northwest Atlantic breeding colonies in the summer of 2022. Results supported all three prediction criteria. Birds in an advanced predefinitive plumage stage were common at colonies (2.0-5.8% of census) even while birds in earlier plumage stages were nearly absent (generally <1% of census). These predefinitive birds were socially engaged while loafing on—and losing fights in—foreign territories. Yet only one out of hundreds of predefinitive birds held a territory or nest. This phenomenon suggests the social conditions of breeding colonies can set the stage for social development that, in turn, sets the stage for life history and plumage evolution.

## INTRODUCTION

Despite growing to full size within months, most seabirds delay reproduction for multiple years (Lack, 1968). The classic explanation for this curious life history is an ecological one: young seabirds get better at foraging as they age (Ashmole, 1963, 1971). If mortality risks decrease as foraging efficiency increases, parents might maximize lifetime reproductive success by delaying reproduction until they pass a foraging efficiency threshold (MacLean, 1986; Pyle et al., 1997). There is especially clear support for this foraging development hypothesis in large, colony-nesting gulls such as Herring Gulls (Larus argentatus). Young Herring Gulls get better at foraging up until their usual age at first reproduction around 4-5 years old (Chabrzyk & Coulson, 1976; Greig et al., 1983; MacLean, 1986).

Yet there is a clue that other developmental processes influence the ecology and evolution of adolescence in these birds: progressive delayed plumage maturation. It takes multiple years for Herring Gulls to advance from the first, brown predefinitive plumage to the gray, black, and white definitive plumages associated with breeding adults (Nisbet et al., 2017). Delayed plumage maturation is common across birds, while multiple, distinct predefinitive plumages are more rare (Hawkins et al., 2012). Distinct stages are best known in gulls (Grant, 1982) along with polygynous lekking birds such as manakins (Schaedler et al., 2021). To be sure, predefinitive plumages can evolve through a broad range of constraints (e.g., limitations in molt evolution; Chu, 1994) or functions (e.g., crypsis to avoid predators; Hawkins et al., 2012). However, it is unclear how foraging efficiency thresholds would relate to gull predefinitive plumages. The former suggests a binary transition between immaturity and maturity, while the latter is a progressive sequence within adolescence itself.

Here, I suggest that an opportunity for social development, rather than foraging development alone, underlies the ecology and evolution of delayed reproduction and delayed plumage maturation in Herring Gulls. Working with Adélie Penguins, Ainley (1975) was the first to seriously suggest that delayed reproduction in seabirds is a function of developing social features such as territories and mates. Indeed, it is well-documented that some seabirds, including Black-legged Kittiwakes (Wooller & Coulson, 1977) and albatross (Pickering, 1989), return to colonies years before breeding. Meanwhile, studies of polygynous, lekking passerines give clues about possible functions of progressive delayed plumage maturation. For Long-tailed Manakins (McDonald, 1993), predefinitive plumages help young males avoid social conflict while they develop the opportunities for, and the content of, sexual displays at leks.

My social development hypothesis proposes that progressive delayed plumage maturation enables colony-related social development (re territories and mates) in monogamous gulls like it enables lek-related social development (re display sites and displays) in polygynous manakins. Testing this hypothesis could require understanding individual behavioral ontogenies, signaling properties of plumage, and subpopulation variation in lifetime reproductive success. A comprehensive study is difficult without tracking cohorts of young gulls across years of low survival (Kentie et al., 2023) and complicated inter-colony movements (Nisbet et al., 2017).

However, there is a fundamental component of the social development hypothesis that I can test with tally counters and rubber boots: do predefinitive gulls have an opportunity for social development at breeding colonies? If delayed reproduction is only a matter of individual development at foraging ground, one would predict that young gulls are rare at colonies—mostly bad foragers, off-colony, not breeding—and only arrive when they are ready to breed. Alternatively, if birds develop central-place foraging skills around the colony itself (c.f., Collet et al., 2020), one would predict that young gulls are common at colonies but have no reason for engagement with territoriality or courtship. There is thus a unique prediction from a social development hypothesis, wherein young gulls enter into the colony for social opportunities themselves (c.f., Ainley, 1975; Ainley et al., 1983). The prediction is that predefinitive birds (A) are present in breeding colonies, (B) are socially engaged, but (C) do not breed.

I tested this prediction at two island breeding colonies of American Herring Gulls (Larus argentatus smithsonianus) in the northwest Atlantic. I asked if predefinitive birds were common at the colony using counts and demographic projections, if they were socially engaged using quantitative tests of attendance and conflict behavior in territorial areas, and if they were breeding using a nest census.

## METHODS

### Study Species

Herring Gulls are large, colony-nesting seabirds in the shorebird radiation Charadriiformes. Most American Herring Gulls (subspecies smithsonianus) migrate from south or central North America to dense summer breeding colonies along Atlantic, Great Lake, or interior lake shores (Nisbet et al., 2017). Established breeders arrive early each spring, often already in pairs, to set up a nesting territory (Tinbergen, 1953). Both parents defend territories, incubate a one-to-three egg clutch, and provision chicks throughout the breeding season (Nisbet et al., 2017). Gulls in my study generally laid eggs in May, incubated through the middle of June, raised chicks through the middle of July, and fledged chicks in August.

Herring Gull territories are social, sexual, and nesting sites, as opposed to foraging sites (Burger, 1984). Foraging sites range from landfills, to fishing boats, to intertidal regions of island colonies (Nisbet et al., 2017; Shlepr et al., 2021). Birds from the same colony can forage from different sites (Enners et al., 2018). For Herring Gulls, the only direct link between territories and foraging involves eating eggs or chicks from other nests (Parsons, 1971; Tinbergen, 1953).

### Plumage stage classification

Herring Gulls molt twice a year, with a complete “basic” molt in the spring and a partial “alternate” molt in the fall (Nisbet et al., 2017). Most Herring Gulls begin breeding at 4 or 5 years old (Chabrzyk & Coulson, 1976; Coulson et al., 1982; Nisbet et al., 2017; Paynter, 1966). These older, breeding birds have iconic “seagull” plumages in the summer: yellow-orange bills with red mandible spots, light eyes with clear pupils, unstreaked white heads and underparts, pale gray mantle, pale gray wings, and black outer primaries with white “mirror” spots (Grant, 1982). A technical name for this plumage is “definitive-cycle alternate” (Wolfe et al., 2010). Here, I just use the term Definitive (Fig. 1).

**Figure 1.**
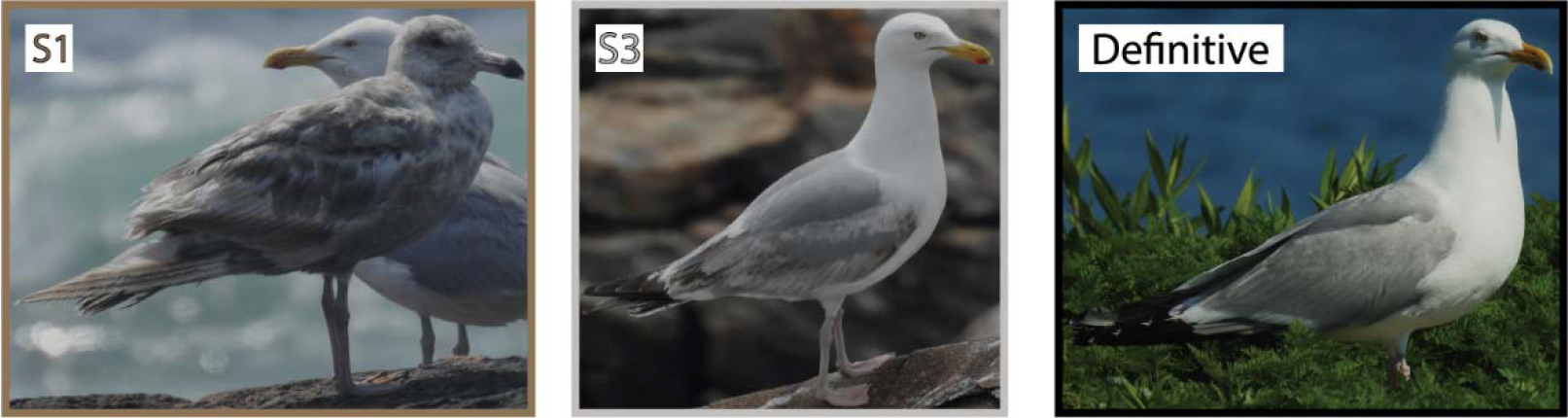
Plumage stage examples for American Herring Gulls (L. a. smithsonianus) in this study. Left: Stage 1 predefinitive, with Definitive behind. Center: Stage 3 predefinitive. Right: Definitive plumage. Photographs from Kent Island and Great Duck Island (July).

Before their definitive plumage, young Herring Gulls pass through a series of progressively bolder predefinitive plumages. I categorized three predefinitive stages for this study (Fig. 1). Classifications were based on Nisbet et al. (2017), public photo repositories (e.g., gull-research.org), and field guide descriptions (e.g., Grant, 1982) of American Herring Gulls:

Stage 1 (S1): Overall-brown plumage, featuring mottled brown feathers with additional gray or black accents across the mantle. Most individuals had head and breast feathers bleached from brown to an inconsistent light gray. All birds in this stage had dark eyes and dark legs. Bills were mostly dark with a lighter base. The guaranteed signs of this stage are shabby, light brown primary flight feathers, which are retained from the juvenile plumage of the previous summer. This stage generally corresponds to first-summer birds (i.e., ∼12 months old, first-cycle formative/alternate plumage).

Stage 2 (S2): Resembling S1 birds except for a distinct gray mantle in contrast to brown upperparts. Compared with S1 birds, S2 birds often had light eyes, lighter legs, and a two-toned bill with a dark tip and broad light base. Head and breast feathers were brown to light gray and were streaked rather than bleached. Tails showed a broad dark band contrasting with a lighter, mottled rump. Wings showed a mix of dark brown secondaries with some patches of mottled brown and gray. Outer primary flight feathers were fresher, dark brown to black, and lacked white mirrors. This stage generally corresponds to second-summer birds (i.e., ∼24 months old, second-cycle alternate plumage).

Stage 3 (S3): Like definitive plumage with four predefinitive distinctions. The first distinction was mottled brown greater and lesser secondary coverts in stark contrast to pale gray median secondary coverts. Second was a dark wingtip, with dark primary coverts and reduced white mirrors. Third was a tripartite bill, with merged red and black spots set on a yellow-orange background. Fourth was a clear and contrasting dark tail band. I occasionally classified an S3 bird even if it was missing one patch (e.g., limited black on the tail, no black on the bill). This stage generally corresponds to third-summer birds (i.e., ∼36 months old, third-cycle alternate plumage).

There are hints that so-called “subadult” birds retain dark markings into their fourth- summer or later. Indeed, I frequently saw birds with one reduced predefinitive plumage patch (e.g., asymmetric dark tail spots, brown tertial spots, or black on the bill). Unlike S3 birds, these birds almost always lacked brown in their secondary coverts. Except for some anecdotal notes on breeding status, I lumped these birds as Definitive.

It occasionally felt difficult to distinguish S1 vs. S2 birds, especially in the fog. Because younger individuals can molt earlier in the summer (Nisbet et al., 2017; Pyle, 2008), classification also became more difficult as the season progressed. I thus lumped the first two plumage stages (S1&2) for all analyses. I probably misclassified some S2 birds as S3 birds, or vice versa. However, the low counts S1&2 birds (see Results: Presence at the colony) suggest my conclusions should withstand occasional errors.

Although my plumage stages putatively correspond to age classes (i.e., S1 birds were ∼12 months old), these stages may be more plastic than generally recognized (B. Buckingham and I. C. T. Nisbet, pers. comm.). Later, I discuss how dynamic relationships between plumage and age would support specific hypotheses about social plasticity.

### Study Sites

I collected count data, quantitative behavioral data, and qualitative behavioral observations at two American Herring Gull breeding colonies on the Atlantic coast of North America (Fig. 1). My main study site was at the Bowdoin Scientific Station on Kent Island, New Brunswick, Canada (44.5828°N, 66.7568°W; Fig. 2). The northern third of this ∼100 ha island is covered by an aging spruce forest, while the southern two-thirds are an open field of tall grasses. American Herring Gulls nest throughout the island, as well as neighboring Sheep and Hay Islands. Kent Island also features ∼20 pairs of Great Black-backed Gulls (Larus marinus). In the 1940’s, Kent Island hosted ∼30,000 breeding gulls (Paynter, 1947). By 2015, the estimated population size had declined to ∼6,000 breeding birds on Kent Island plus ∼3,000 breeding birds on Sheep and Hay Islands (Bennett et al., 2017). Kent Island is thus a low-density and declining colony.

**Figure 2.**
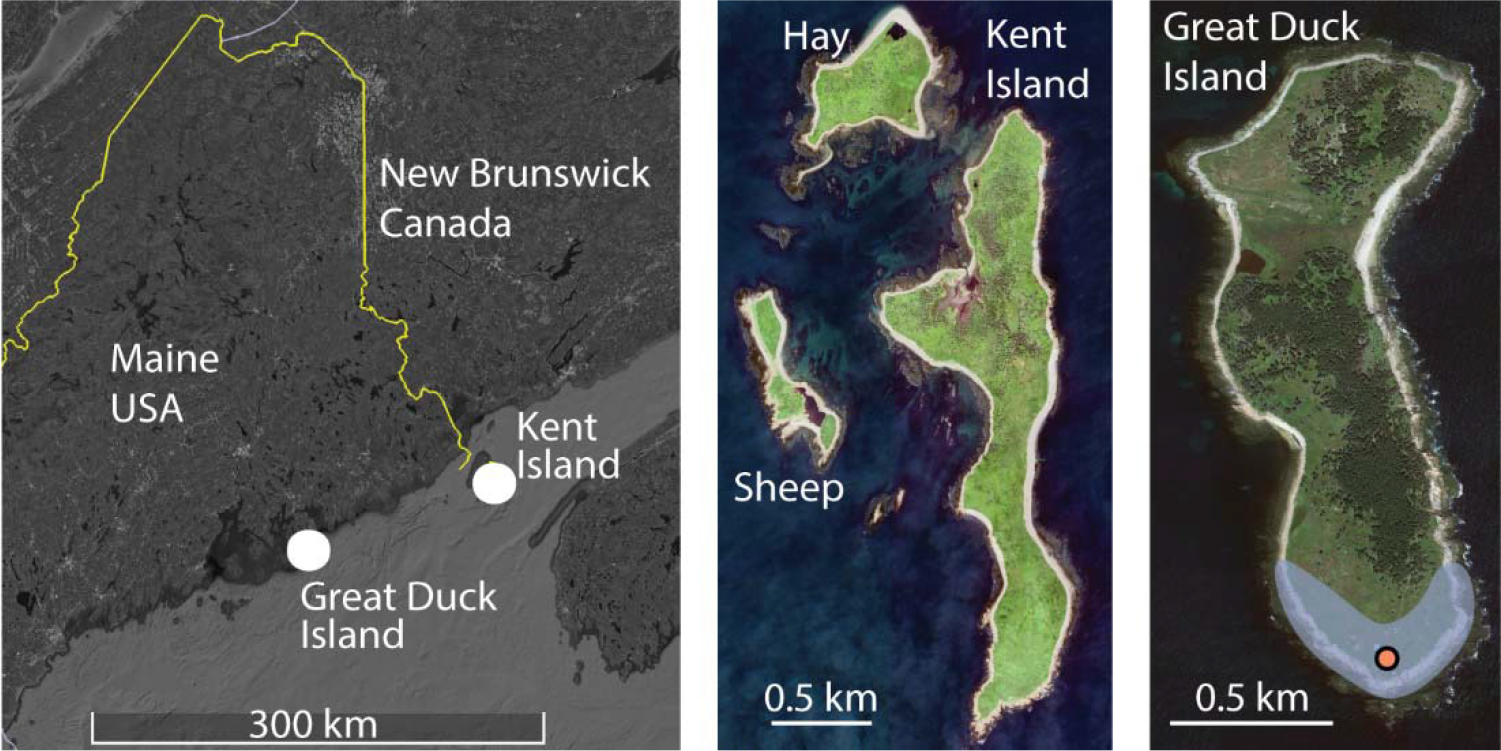
American Herring Gull breeding colonies surveyed for this study. Left: Regional map. Center: Kent Island off Grand Manan, New Brunswick, Canada. Right: Great Duck Island off Bar Harbor, Maine, USA. The main colony on Great Duck Island is highlighted in blue with a dot marking the lighthouse observation tower. Satellite images from Google Earth.

I made additional observations at the Alice Eno Field Station on Great Duck Island, Maine, USA (44.1449°N, 68.2458°W; Fig. 2). The American Herring Gulls across Great Duck Island have increased from ∼600 birds in 1985 to ∼1300 birds in 2021 (J. Anderson pers. comm.), making it a smaller but denser colony than Kent Island. The main breeding colony on the south shore features a lighthouse observation tower.

### Ethical Note

All methods were approved by the Bowdoin Scientific Station and Alice Eno Field Station and waived for additional approval by Yale University and Bowdoin College.

### Presence at the colony

I conducted hundreds of walking counts to estimate the plumage stage distribution of American Herring Gulls at summer breeding colonies (June - August, 2022). I counted S1, S2, S3, and total birds, inferring Definitive counts as total minus S1-3 counts. I combined S1 and S2 counts for all analyses. All counts excluded young-of-year. My counts are not equivalent to historical or regional demographic surveys because they include both breeding and non-breeding birds. I used repeated counts to assess the impact of observation error (Supplementary Material). All data wrangling and analyses were done with the tidyverse packages in R (R Core Team, 2021; Wickham et al., 2019).

I divided Kent Island into 10 census areas (Fig. 3). I counted four of those areas (East Beach, West Beach, South Basin, and Downtown) one to three times per day for much of the breeding season (June 1 - July 7, July 23 - August 15, 2022). I counted the entire island once per week during this period. I surveyed Hay Island twice (June 21 and August 7, 2022). I counted gulls by walking down the rocky beach, tallying every bird I passed and discounting birds flying in from behind me. On Great Duck Island, I did 10 counts of the main breeding colony from the observation tower (July 14-20, 2022).

**Figure 3.**
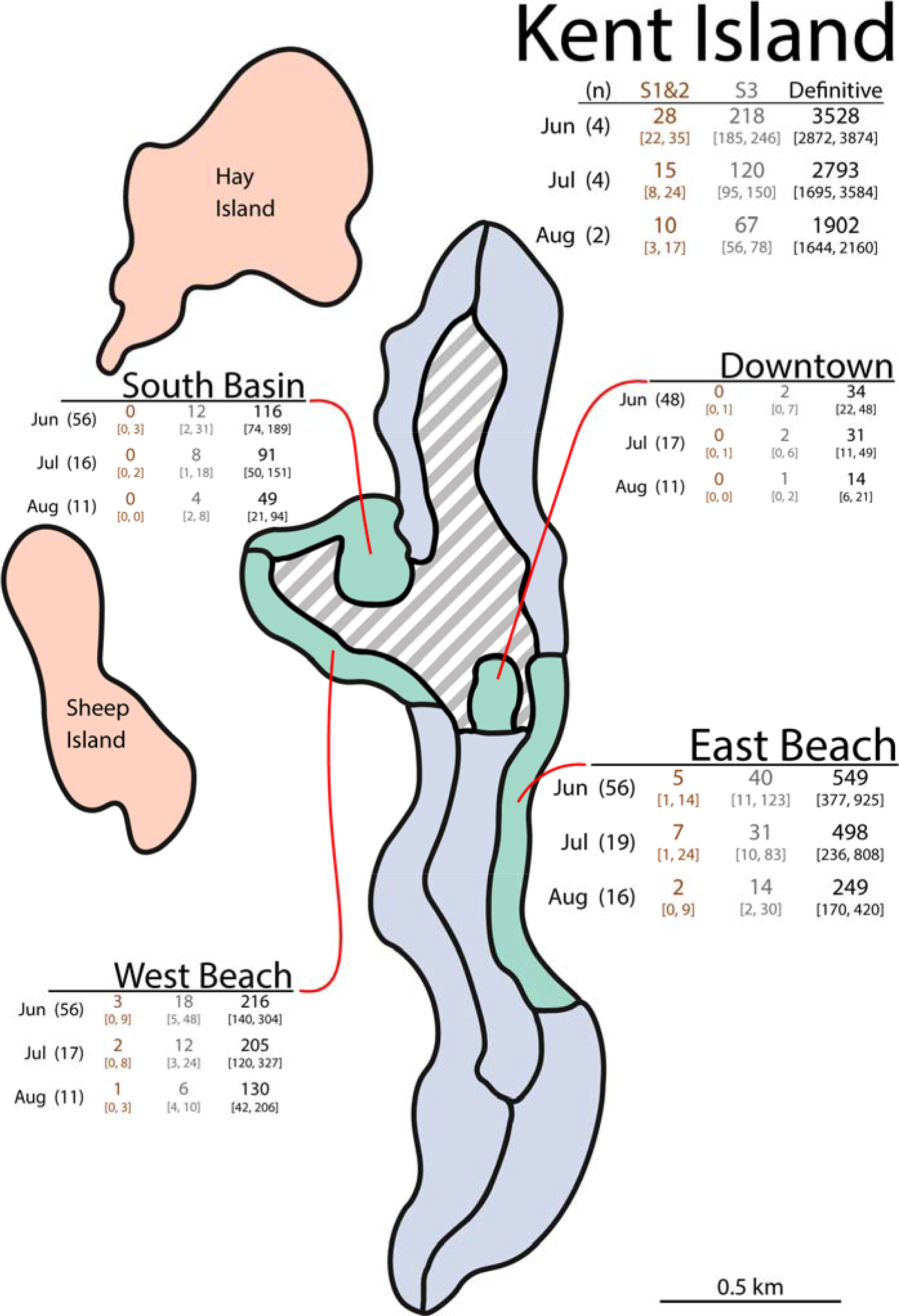
Census areas on Kent Island, New Brunswick, Canada. I counted American Herring Gulls on the four green segments at least once a day for much of the 2022 breeding season (June 1-July 7 and July 23 - August 15, 2022). I counted the entire island, including blue segments, once a week during this period. I did not formally survey the bogs, fields, and forest in the dashed gray region. Charts show count summaries from each of the main areas. Counts are given as mean [min, max] separated by plumage class (predefinitive S1&2, predefinitive S3, and Definitive) across (n) counts per month.

In 2021, I conducted one island-wide count of Kent Island (August 17-18, 2021) and 15 lighthouse counts of Great Duck Island (July 1-6, 2021). I was only equipped to distinguish predefinitive (S1-3) vs. Definitive birds during 2021 counts.

### Demographic expectations

I compared observed plumage class distributions to expected age class distributions using demographic parameter estimates from Kent Island. The expected stable age distribution of a population is given by the right eigenvector of the Leslie matrix of reproduction and survival parameters, while population growth rate is given by a positive real eigenvalue (Caswell, 2001). I first calculated the expected stable age distribution by taking rough historical estimates of per-capita annual fledged offspring (0.92), yearling survival rate (0.56), annual adult survival rate after the first year (0.82), and an age at reproduction of 4 years (Freeman & Morgan, 1992; Paynter, 1966). Consistent with historical records, these values correspond to an increasing population growth rate ∼1.059.

However, recent Kent Island censuses show a decreasing gull population with annual growth rate ∼0.953 (from 5,926 breeding pairs in 2001 to 3,004 breeding pairs in 2015; Bennett et al., 2017). The impact of population growth rate differences on expected age distributions depends on which demographic rates have changed. The proportion of young birds at stable age distribution would be higher given shrinking adult survival. Alternatively, the proportion of young birds would be lower given shrinking yearling survival. I thus calculated bounds on expected young bird proportions by simulating changes to survival parameters until population growth rate declined from 1.059 (historical estimate) to 0.953 (modern estimate). The upper estimate on young bird proportions comes from iteratively decreasing adult survival rate while the lower estimate comes from iteratively decreasing yearling survival rate.

### Social engagement

Even if young birds are present on a breeding island, one possibility is that they are merely loafing in isolated regions such as the intertidal. I thus quantified social engagement by asking whether young gulls were present and engaged among the actual breeding territories of colonies. Statistical tests were supplemented with qualitative behavioral observations on the territoriality, conflict, and courtship throughout the summer of 2022 (Supplementary Material).

On Kent Island, I quantified social engagement by separately counting the territorial and non-territorial areas of East Beach. The territorial area (featuring all territories and nests) fell above the midline of East Beach and onto the vegetated berm. The non-territorial area (featuring zero nests or territories) fell below the midline of the beach and out to the rocky intertidal. I counted the birds in the territorial and non-territorial areas in the reverse direction (northbound) of my usual (southbound) East Beach counts (June 7 - July 4, 2022). It was tricky to assess both plumage and location of every bird while walking down the beach. As a result, I ran 24 counts of all birds (lumping S1-3 plus Definitive) and 20 counts of just S1-3 birds (excluding Definitive). I compared the proportion of birds of different ages classes (S1&2 vs. S3 vs. all birds) in the territorial area using a Kruskal-Wallis rank sum test followed by three pairwise asymptotic Wilcoxon-Mann-Whitney tests (mid-rank ties; Bonferroni-corrected α = 0.016; coin::wilcox_test: Zeileis et al., 2008).

On Great Duck Island, I was able to monitor individual behaviors by tracking flying birds from the lighthouse observation deck (July 14-20, 2022). I haphazardly picked an S3 or Definitive bird flying above the height of the keeper’s cottage. I recorded whether that bird landed in a territorial area (i.e., places on the beach, berm, or meadow closely associated with nests, chicks, or breeding adults) or a non-territorial area (i.e., places not associated with nests, including station buildings, intertidal, or empty patches of meadow). Flights where the focal bird flew into the forest or off to sea were excluded. Across 457 individual flights, I tested whether S3 birds were less likely than Definitive birds to land in territorial areas using χ^2^ tests.

I also used Great Duck Island flights to investigate pairwise displacement conflicts. Displacement conflicts occurred when the landing bird (whether S3 or Definitive) touched down near a sitting bird (whether S3 or Definitive) and one individual was quickly forced off the site by the other. I marked the winner as the bird that remained at the landing site after a conflict. This conflict dataset included 457 tracked flights, 107 of which ended in conflict, plus 30 focal observations of sitting birds. I first asked whether S3 flights were more likely to end in conflict than Definitive flights using χ2 tests. After including focal observations of sitting birds, I then asked whether S3 birds lost more conflicts than Definitive birds using Fisher’s exact tests. Some of these datapoints were pseudoreplicates (Hurlbert, 1984). There were relatively few S3 birds in the colony each day and the same Definitive birds took multiple trips around the colony. However, I attempted to observe every different S3 bird at the colony each day and tried to avoid resampling Definitive birds nesting close to the lighthouse.

### Breeding status

I surveyed nests, chick-groups, and attending parents across four segments of Kent Island (East Beach, West Beach, South Basin, and Downtown; Fig. 3). I made one nest count on June 7, 2021. Later, I made two replicate counts of nests and chick-groups on July 4 and July 5, 2022. I recorded whether nests or chicks were being actively attended by at least one predefinitive parent.

## RESULTS

### Presence at the breeding colony

Full-island counts featured a few dozen S1&2 birds, a few hundred S3 birds, and a few thousand Definitive birds (Figs. 3-4). In June 2022 on Kent Island, I counted an average of 0.8% S1&2 birds, 5.8% S3 birds, and 93.4% Definitive birds. Relative proportions of S3 birds declined throughout the season. By August, S3 birds were 3.5% of total (Table 1). One June count of Hay Island featured 3 S1&2 birds (0.4%), 20 S3 birds (2.5%), and 782 Definitive birds (97.1%), while one August count featured 1 S1&2 bird (0.2%), 19 S3 birds (4.4%), and 408 Definitive birds (95.4%).

**Figure 4.**
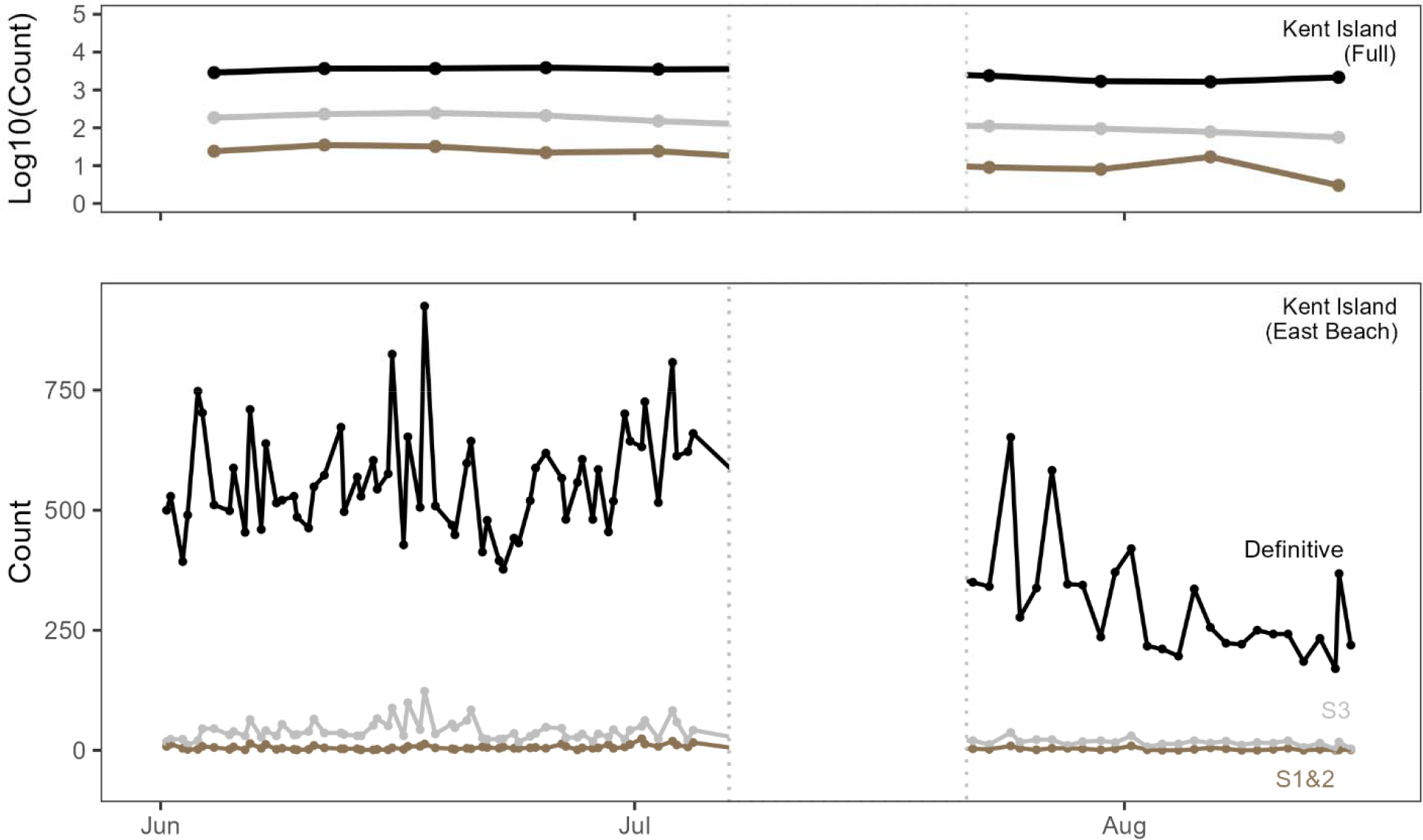
Plumage class counts of American Herring Gulls on Kent Island, summer 2022 (brown = S1&2 predefinitive, gray= S3 predefinitive, black = Definitive). Top: Log-scale counts from weekly full-island surveys. Bottom: More frequent counts from the East Beach area.

**Table 1.**
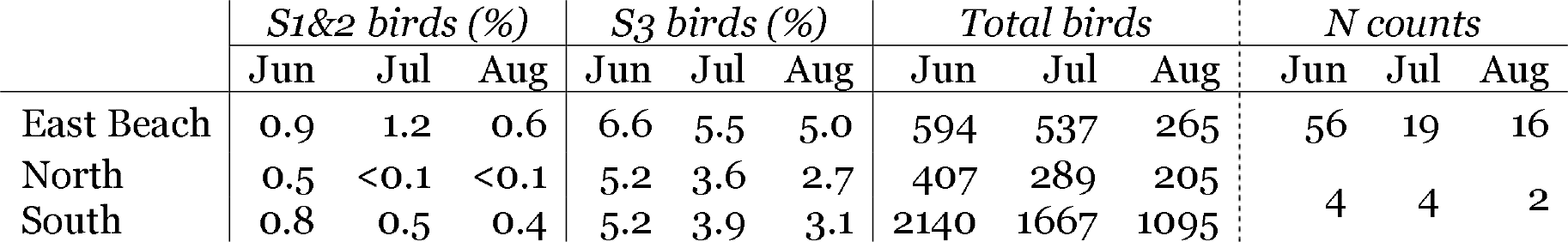
Representation of predefinitive American Herring Gulls across Kent Island. “North” combines the lower density North West and North East areas of the island (see Fig. 3). “South” combines the higher density South West and South East segments. All values averaged across N counts during summer 2022.

More frequent counts of East Beach, Kent Island showed similar plumage class distributions (Figs. 3-4, Table 1). East Beach counts varied widely throughout the season (mean = 524±180 birds, range = 172-1,061 birds, n = 91 counts; Fig. 4). Counts swung by hundreds of birds even within an individual day, suggesting that both breeding and nonbreeding birds were moving into, around, and out of the colony with the sun and tide. The maximum count of total, Definitive, and S3 birds occurred late in the afternoon of June 16, featuring 925 Definitive and 123 S3 birds. However, the maximum count of S1&2 birds was in the morning of July 1 (Canada Day), with 24 S1&2 birds (3.4% count total).

Plumage class distributions were generally consistent across other areas of Kent Island (Fig. 3). On West Beach (47-350 birds, n = 84 counts), S3 birds were 6.7% and S1&2 birds were 1.0% of totals across the 2022 season. At South Basin (23-223 birds, n = 83 counts), S3 birds were 8.7% and S1&2 birds 0.1% of total. The small Downtown section (6-53 birds, n = 76 counts) at the center of the field station had fewer predefinitive birds, with S3 birds at 4.2% of total and only two S1&2 birds ever spotted.

Along the coast of Kent Island, gulls were generally sparser in the north and denser in the south. Despite these differences, representation of S3 birds was nearly identical across northern and southern areas. East Beach was of intermediate overall density but featured an equal, or greater, representation of predefinitive birds (Table 1).

Great Duck Island had a lower proportion of predefinitive birds than Kent Island. Counts during one week in July, 2022 (412-569 birds, n = 10 counts) featured an average 2.0% (range 6-18) S3 birds and 0.5% (1-6) S1&2 birds.

Preliminary surveys from 2021 were consistent with 2022 plumage class distributions. A single full-island count of Kent Island in August 2021, gave 3,595 total birds with 3.1% predefinitive birds. Counts of the main Great Duck Island colony during one week in July 2021 (392-856 birds, n = 15 counts) featured an average of 4.4% (range 7-62) predefinitive birds. Although I did not confidently distinguish S1&2 vs. S3 birds during these preliminary counts, my notes suggest that most predefinitive birds in 2021 were S3.

A limited outbreak of highly pathogenic avian influenza ran through Kent, Sheep, and Hay Islands starting on June 27, 2022 (Caliendo et al., 2022). This outbreak resulted in the deaths of dozens of birds, the details of which will be reported elsewhere. Nearly all the influenza-related carcasses found on Kent Island were Definitive birds, suggesting the outbreak should not impact my conclusions. Given limited information on normal phenology, however, some seasonal shifts in counts or proportions could have been altered by the outbreak.

### Demographic expectations

Using the rough historical estimates for Kent Island gulls, the stable age distribution predicts 40% year one and two (Y1&2) birds, 14% Y3 birds, and 46% Y4+ birds. A reduction in yearling survival rate from 0.56 to 0.23 drops estimated population growth rate from 1.059 to 0.953. The result is an expected distribution of 26% Y1&2, 10% Y3, and 64% Y4+ birds. A decrease in adult survival rate from 0.82 to 0.73 also drops expected population growth rate to 0.953 but barely shifts the expected distribution (42% Y1&2, 14% Y3, 44% Y4+). Given that I observed few S1&2 birds, I calculated the relative proportions of Y3 and Y4+ birds. Excluding Y1&2 birds shifted the expectation range to 14-24% Y3 birds.

### Social engagement

Gulls in S3 plumages on East Beach, Kent Island showed distinct, transitional associations with territorial areas (Fig. 5). The proportion of S3 birds in the territorial area (0.31) was significantly smaller than the proportion of total birds in the territorial area (0.66) but significantly larger than proportions from a handful of S1&2 birds (0.01; overall Kruskal-Wallis rank sum test, H = 56.36, P < 0.001; all pairwise Asymptotic Wilcoxon-Mann-Whitney P < 0.001).

**Figure 5.**
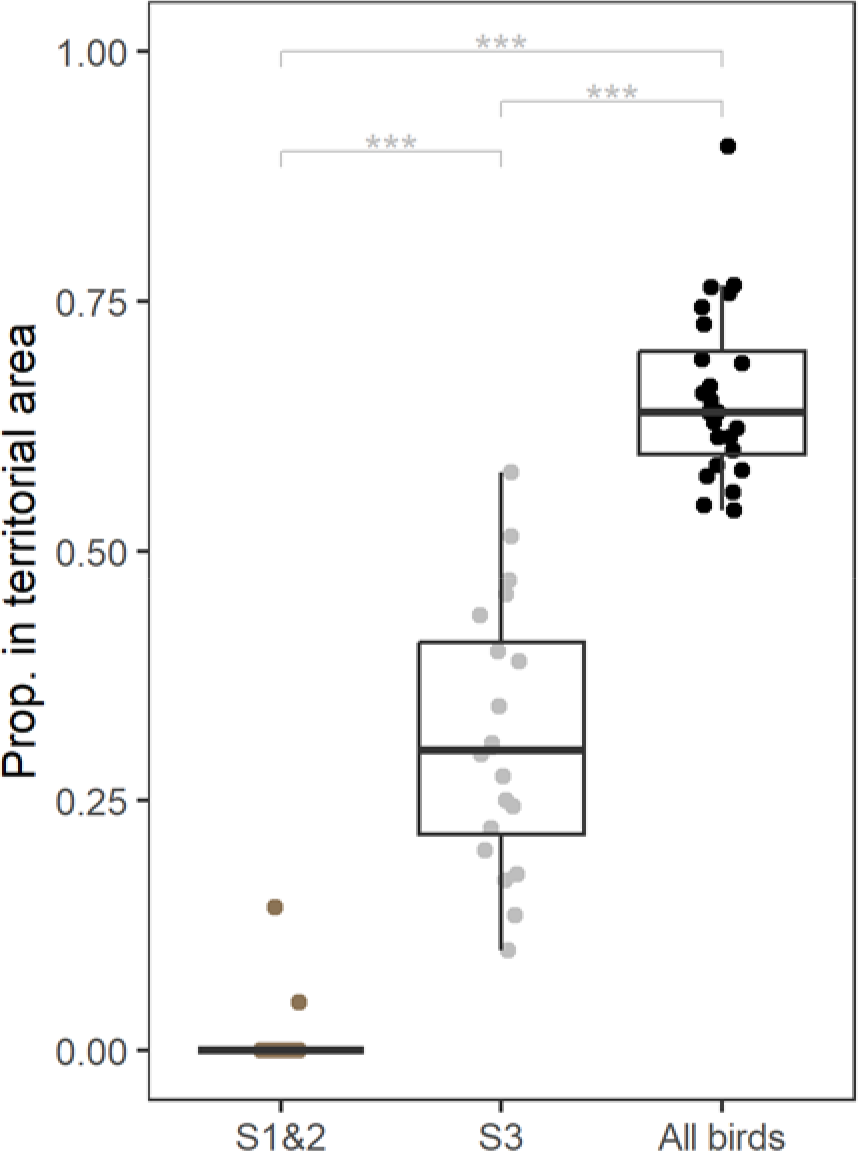
Proportion of American Herring Gulls in territorial areas on East Beach, Kent Island, separated by plumage class (predefinitive S1&2, predefinitive S3, all birds). “All birds” counts were conducted separately from S1-3 counts and represent substantially higher raw values (S1&2: 7±6 birds, 20 counts; S3: 51±26 birds, 20 counts; All birds: 636±83 birds, 24 counts). Asterisks (***) indicate P < 0.001 for pairwise Wilcoxon-Mann-Whitney tests of proportions.

Predefinitive birds on Great Duck Island also showed transitional associations with territorial areas. By tracking individual flights, I saw S3 birds make significantly fewer landings in territorial areas than Definitive birds (S3: 63/122 landings territorial; Definitive: 228/335 landings territorial; χ^2^ = 9.73, P = 0.002; Table 2) but significantly more than a zero expectation (S3: 63/122 landings territorial; expected: 0/122 landings territorial; χ^2^ = 82.25, P < 0.001).

**Table 2.** Landing sites of American Herring Gulls in predefinitive (S3) or Definitive on Great Duck Island. Individual birds touched down on either territorial or non-territorial areas, at which point they either landed without disturbance or engaged in a displacement conflict. S3 birds were significantly less likely to land in territorial areas than Definitive birds, but did so significantly more than a zero-expectation. S3 birds were also significantly more likely to get into conflicts when landing in territorial areas.

| *Landing bird* | *Territorial landing* |  | *Non-territorial landing* |  |
| --- | --- | --- | --- | --- |
|  | *Conflict* | *No conflict* | *Conflict* | *No conflict* |
| S3 | 29 | 34 | 11 | 48 |
| Definitive | 49 | 179 | 18 | 89 |

These predefinitive birds got into significantly more conflicts than Definitive birds when landing in territorial areas (χ^2^ = 13.93, P < 0.001; Table 2). On the other hand, S3 and Definitive birds were equally likely to get into conflicts when landing in non-territorial areas (Table 3; χ^2^ = 0.01, P = 0.93; Table 2). Finally, S3 birds were significantly less likely than Definitive birds to “win” conflicts as either the landing or the sitting bird in both territorial and non-territorial areas (Table 3; S3 landing bird vs. Definitive landing bird, wins across all sitting birds: territorial odds ratio = 0.08, non-territorial odds ratio = 0.06; S3 sitting bird vs. Definitive sitting bird, wins across all landing birds: territorial odds ratio = 0, non-territorial odds ratio = 0.08, P = 0.011; Odds ratios from Fisher’s exact tests, P < 0.001 unless noted). Qualitative observations helped clarify that S3 birds in territorial areas did not actually establish or maintain territories (Supplementary Material).

**Table 3.** Conflict win:loss ratios among predefinitive (S3) and Definitive American Herring Gulls on Great Duck Island. Wins given in terms of the landing bird. Events include the conflict landings from Table 2 plus 30 focal observations of sitting birds. Although sample sizes are small, S3 birds were significantly more likely than Definitive birds to lose conflicts in both territorial and non-territorial areas.

| *Landing bird* | *Sitting bird* |  |  |  |
| --- | --- | --- | --- | --- |
|  | *Territorial area* |  | *Non-territorial area* |  |
|  | S3 | Definitive | S3 | Definitive |
| S3 | 1:0 | 1:28 | 0:0 | 2:9 |
| Definitive | 19:0 | 15:35 | 12:1 | 10:4 |

### Breeding status

On June 7, 2022, I counted 479 active nests across the East Beach, West Beach, South Basin, and Downtown segments of Kent Island. During this initial count, I noted 8 nests (1.7%) with any, maximally liberal, association with a predefinitive bird. However, only one nest (0.2%) had an attending parent in S3 predefinitive plumage (see Supplementary Material for details).

Two subsequent counts of the same areas—including both nests and hatched chick groups—indicated 201 (July 4) and 218 (July 5) clutches. Once again, just the one clutch was attended by the same S3 bird. At no point did I see another predefinitive parent on either Kent Island or Great Duck Island. Anecdotally, it was not uncommon to see a nest with a “subadult” parent (see Methods: Plumage stage classification).

## DISCUSSION

American Herring Gulls in a specific predefinitive plumage stage (S3) were present and socially engaged, but not breeding, at summer breeding colonies on Kent Island, Canada and Great Duck Island, USA. These results are consistent with the hypothesis that young gulls in predefinitive plumages have an opportunity for social development at the breeding colony.

In terms of presence at the breeding colony, S3 gulls were frequent. In June 2022 on Kent Island, these young birds made up an average of 6.6% of East Beach counts and 5.8% of island-wide counts (Figs. 3-4, Table 1). There were fewer S3 birds (2.0%) at the smaller Great Duck Island colony in July. The proportions of predefinitive birds in 2022 were consistent with preliminary 2021 counts at both colonies, even though these counts presumably correspond to different birds passing through plumage stages.

How many predefinitive birds should we expect at colonies? Without knowing how individual attendance patterns across cohorts of birds articulate with—or historically articulated with—processes of selection, there is no threshold value that could support or reject a life history hypothesis. [Readers may attempt the exercise and contact me if they determine a value >0%.] However, rough demographic projections offer a rough guide. If S3 birds are all, and the only, three-year-old birds, old demographic values from Kent Island projected 14-24% S3 representation. In June on Kent Island, I might expect 906 S3 birds (24% of 3,775 birds; Fig. 3) when I only observed an average of 218 S3 birds (24% expected). This discrepancy could be understood in several ways, none of which can be levied in support or rejection of a life history hypothesis. First, different S3 individuals could each attend the colony for short periods of time. Observations of banded birds would both clarify individual activity and help pinpoint why proportions of predefinitive birds declined throughout the season (Fig. 4, Table 1). Second, additional S3 birds may attend other colonies. Young Herring Gulls do not always return to their natal colony (Chabrzyk & Coulson, 1976; Nisbet et al., 2017). Satellite tracking technology should finally help us understand these movements in greater detail (e.g., Frankish et al., 2022). Third, a subset of birds may not attend a colony at all in S3 plumage. Chabrzyk and Coulson (1976) show that recruitment age is younger in low-density Herring Gull colonies, meaning pre-breeding dynamics could themselves vary with ecological and social contexts. Indeed, the lower-density, declining Kent Island colony in this study had a higher proportion of S3 birds (4.3% in July 2022) than the denser, increasing Great Duck Island colony (2.0% in July 2022). Similar comparisons across years and colonies are needed to track the metapopulation dynamics of young gulls.

Rather than using rough demographic projections, my data provide far more direct insight via internal comparison between S1&2 vs. S3 birds. While there were hundreds of S3 birds on Kent Island, there were only a handful of S1&2 birds (<1% of counts; Figs. 3-4, Table 1). These birds are missing from breeding colonies even though any stable demographic scenario requires more S1 birds than S3 birds to be alive. Either gull populations have faced a new precipitous crash, or older adolescent birds uniquely return to the breeding colony at much higher rates than younger birds.

In terms of social engagement, predefinitive S3 gulls frequently loafed on or landed in territorial areas (Fig. 5, Table 2). Qualitative observations confirmed that these birds were not actually holding territories (Supplementary Material). Indeed, S3 birds were significantly more likely than Definitive birds to face a conflict when landing in a territorial area (Table 2) and were significantly more likely to be displaced in both territorial and non-territorial areas (Table 3).

In terms of breeding status, just one (0.2%) of the nests censused on Kent Island was attended by an S3 parent. Predefinitive birds were not breeding despite having a suitable vocal repertoire and occasionally engaging in incomplete courtship (Supplementary Material). This result confirms previous statements that predefinitive Herring Gulls rarely breed (Coulson et al., 1982; Tinbergen, 1953).

If individual foraging skill was all that mattered to young gulls, then social engagement at the breeding colony should align with attempted reproduction. Capable foragers would have no special difficulties establishing territories, finding and maintaining mates, and attempting reproduction (regardless of experience-related differences in breeding outcome; Curio, 1983; Pyle et al., 1997). American Herring Gulls do something different. Birds in a special plumage stage (S3) occupy the colony while neither breeding nor attempting to breed.

An alternate hypothesis is that young seabirds returning to a breeding colony are prospecting for local foraging conditions (Ashmole, 1963; Lack, 1954). Prospecting is unlikely to work for American Herring Gulls, whose foraging grounds and strategies differ widely within colonies and have little to do with territorial conditions (Nisbet et al., 2017). It is also conceivable that young birds could arrive at the colony to develop colony-specific foraging skills (c.f., Collet et al., 2020). This hypothesis is insufficient for two observations: (A) S3 birds were socially engaged and (B) S3 birds were present but S1&2 birds were mostly absent. If foraging development at the colony was the only relevant factor for S3 birds, I would not predict these birds to risk their necks in territorial areas. Alternatively, if foraging development at the colony was relevant for all young birds, then S1&2 birds would be frequent at colonies. My results are consistent with social development and neither consistent with nor predicted by foraging development alone.

Qualitative behavior differences (off-colony vs. on-colony) across qualitative plumage classes (S1&2 vs. S3) suggest American Herring Gulls pass through multiple developmental phases before breeding. Specifically, young gulls may go through a period of foraging development off the colony followed by a period of social development on the colony. There is good evidence that gulls become more efficient foragers as they age (Greig et al., 1983; MacLean, 1986). There are also qualitative differences in foraging movements for the youngest birds. For example, Paynter (1947) found most Kent Island gulls taken by fishermen were first- or second-year birds. My own peak count of S1&2 birds from East Beach, Kent Island was the morning of July 1. This is the date the local lobster fishery closes for the season, hinting that the youngest gulls were moving in response to foraging opportunities. Ainley et al. (1983) propose the same developmental phases—foraging, then social—for Adélie Penguins, and we have direct data from individually-tracked Wandering Albatross. Young albatross forage differently at first, but they quickly catch up to older birds and still do not breed for several years (Riotte-Lambert & Weimerskirch, 2013).

What does a social development phase mean for young seabirds? One simple suggestion is that young gulls do not breed because there is a survival cost to reproduction (Pyle et al., 1997; Wooller & Coulson, 1977). Another simple suggestion might involve alternative reproductive tactics such as extra-pair copulation (c.f., Taborsky et al., 2008). There is indeed evidence of courtship and copulation by young Herring Gulls (Tinbergen, 1953; Supplementary Material). However, female Herring Gulls delay both reproduction and plumage without dumping eggs in foreign nests (Nisbet et al., 2017). Further, hundreds of predefinitive birds persisted on Kent Island through July and August, long after any successful nest is established (Fig. 4). In any case, reasoning based on reproductive costs or alternative strategies offer only tautological explanations for age-related polymorphisms such as delayed reproduction. If there are costs to reproduction, I would need to say why costs differ between young and old birds. If there are alternate reproductive strategies, I would need to say why strategies differ between young and old birds.

In other words, I would need to say something about development. Ainley et al. (1983) suggest that adolescent penguins need rookery experience to eventually obtain territories or mates. Bradley and Wooller (1990) make general references to “social environments” and “conventions.” More explicit is Pickering (1989), who documents the ways in which Wandering Albatross begin courting multiple partners several years before breeding. Pickering (1989) suggests young albatross must establish and maintain a pair-bond before reproduction. This is a radical suggestion, couched as it is in the subtle terms of seabird biology. It is a material fact that albatross require biparental care to raise young (Tickell, 1968). Developing a pair-bond is thus a material requirement of reproduction, like feathers or gonads. We know plenty about feathers and gonads. We have barely reckoned with what it means for a bird to develop a pair-bond, let alone grapple with a social convention.

My observations suggest that social development in American Herring Gulls has something to do with the social capacity for territoriality. This capacity is different than obtaining a territory per se (Coulson, 2001). There was plenty of room for nests on Kent Island, yet the sparsest areas in the north had no glut of predefinitive birds (Table 1). Meanwhile, predefinitive birds were getting into—and losing—fights in foreign territories (Table 2, Supplementary Material). In his discussion of egg thieves, Tinbergen (1953) suggests that young gulls may operate outside of the social conventions of territoriality itself. I did not see egg thieves (Supplementary Material), but the broader idea fits. Young gulls might develop with respect to territorial conventions (“what does lunging at another bird do?”) even if they are neither developing territories (“where is the border at which I lunge?”) nor learning per se (“what is a lunge?”).

My data suggest that the social development of territoriality sets the ecological stage for predefinitive plumage evolution in gulls. Studies from tropical lekking birds show how predefinitive plumage stages can enable sociosexual development. For example, the males of several manakin species also pass through multiple predefinitive plumage stages (Schaedler et al., 2021). In Long-tailed Manakins, predefinitive plumages help mitigate aggression while young males navigate cooperative social hierarchies needed for sexual displays (McDonald, 1993). In this study, American Herring Gulls returned to the colony in a distinct plumage stage (S3) that simultaneously signals both youth (e.g., brown wing coverts, black tail band) and maturity (i.e., white head, silver scapulars). This plumage is consistent with the need for young gulls to mediate social engagement before breeding. Distinct developmental phases—foraging development off the colony followed by social development on the colony—are further consistent with the distinct predefinitive plumage stages. We might now investigate how foraging and social environments covary among those seabirds with delayed plumage maturation (e.g., Common Terns, Wandering Albatross; Bridge & Nisbet, 2004; Tickell, 1968) and those without (e.g., Adélie Penguins, Leach’s Storm-Petrels; Ainley et al., 1983; Pollet et al., 2021).

We might also learn more about the ontogeny of seabird plumages. For example, I found that American Herring Gulls are socially engaged at the colony in the S3 plumage. I assumed that predefinitive plumage stages—S1, S2, S3—corresponded to age classes—12 months, 24 months, 36 months. Yet banding records show variations such as S3-like plumages before the third year (B. Buckingham pers. comm.). Such plastic delayed plumage maturation has been studied in some cases (especially Red-backed Fairy-wrens: Welklin et al., 2021) and suggested in others (manakins: Schaedler et al., 2021). If individuals vary in the age at which they advance to S3 plumage, this suggests variation in the competency to begin the social phase of development. Birds may also differ by the number of years spent in S3 plumage. This pattern would suggest variation in the process of social development itself. Gull plumage thus raises questions about not only about ecology and behavior, but also development and plasticity. From sexual bimaturism in Orangutans (Kralick et al., In review) to sexual transition in wrasses (Lowe et al., 2021), we are just beginning to map the dynamics of socially-mediated life histories. It seems that gulls—once premier models of social behavior (Tinbergen, 1953)—still have a lot to teach us.

## ACKNOWLEDGEMENTS

I am deeply grateful to the staff, students, and faculty on Kent Island and Great Duck Island, especially Patricia Jones, Ian Kyle, Wriley Hodge, and John Anderson. I thank Richard Prum, Stephen Stearns, Bruce Buckingham, Ian Nisbet, and Kate Shlepr for comments on the topic or the manuscript. This work was supported by NSF GRFP (#DGE1752134) and the W. R. Coe fund from Yale University.

## REFERENCES

Ainley, D. G. (1975). Development of reproductive maturity in Adélie Penguins. In The biology of penguins (pp. 139–157). Macmillan.

Ainley, D. G., LeResche, R. E., & Sladen, W. J. (1983). Breeding Biology of the Adélie Penguin. University of California Press.

Ashmole, N. P. (1963). The regulation of numbers of tropical oceanic birds. Ibis, 103b(3), 458– 473.

Ashmole, N. P. (1971). Sea bird ecology and the marine environment. In D. S. Farner, J. R. King, & K. C. Parkes (Eds.), Avian Biology (Vol. 1, pp. 223–286). Academic Press, New York.

Bennett, J., Jamieson, E., Ronconi, R., & Wong, S. (2017). Variability in egg size and population declines of Herring Gulls in relation to fisheries and climate conditions. Avian Conservation and Ecology, 12(2).

Bradley, J. S., & Wooller, R. D. (1990). Philopatry and Age of First-breeding in Long-lived Birds. Acta XX Congressus Internationalis Ornithologici, 3, 1657–1665.

Bridge, E. S., & Nisbet, I. C. T. (2004). Wing Molt and Assortative Mating in Common Terns: A Test of the Molt-Signaling Hypothesis. The Condor, 106(2), 336–343.

Burger, J. (1984). Pattern, mechanism, and adaptive significance of territoriality in Herring Gulls (Larus argentatus). Ornithological Monographs, 34, iii–92.

Caliendo, V., Lewis, N. S., Pohlmann, A., Baillie, S. R., Banyard, A. C., Beer, M., Brown, I. H., Fouchier, R. a. M., Hansen, R. D. E., Lameris, T. K., Lang, A. S., Laurendeau, S., Lung, O., Robertson, G., van der Jeugd, H., Alkie, T. N., Thorup, K., van Toor, M. L., Waldenström, J., … Berhane, Y. (2022). Transatlantic spread of highly pathogenic avian influenza H5N1 by wild birds from Europe to North America in 2021. Scientific Reports, 12(1), Article 1.

Caswell, H. (2001). Matrix population models. John Wiley & Sons, Ltd.

Chabrzyk, G., & Coulson, J. C. (1976). Survival and Recruitment in the Herring Gull Larus argentatus. Journal of Animal Ecology, 45(1), 187–203.

Chu, P. C. (1994). Historical examination of delayed plumage maturation in the shorebirds (Aves: Charadriiformes). Evolution, 48(2), 327–350.

Collet, J., Prudor, A., Corbeau, A., Mendez, L., & Weimerskirch, H. (2020). First explorations: Ontogeny of central place foraging directions in two tropical seabirds. Behavioral Ecology, 31(3), 815–825.

Coulson, J. C. (2001). Colonial breeding in seabirds. In Biology of marine birds (pp. 100–127). CRC Press.

Coulson, J. C., Duncan, N., & Thomas, C. (1982). Changes in the Breeding Biology of the Herring Gull (Larus argentatus) Induced by Reduction in the Size and Density of the Colony. Journal of Animal Ecology, 51(3), 739–756.

Curio, E. (1983). Why do young birds reproduce less well? Ibis, 125(3), 400–404.

Enners, L., Schwemmer, P., Corman, A.-M., Voigt, C. C., & Garthe, S. (2018). Intercolony variations in movement patterns and foraging behaviors among herring gulls (Larus argentatus) breeding in the eastern Wadden Sea. Ecology and Evolution, 8(15), 7529–7542.

Frankish, C. K., Manica, A., Clay, T. A., Wood, A. G., & Phillips, R. A. (2022). Ontogeny of movement patterns and habitat selection in juvenile albatrosses. Oikos, 2022(6), e09057.

Freeman, S. N., & Morgan, B. J. T. (1992). A Modelling Strategy for Recovery Data from Birds Ringed as Nestlings. Biometrics, 48(1), 217–235.

Grant, P. J. (1982). Gulls: A guide to identification. Buteo Books.

Greig, S. A., Coulson, J. C., & Monaghan, P. (1983). Age-related differences in foraging success in the herring gull (Larus argentatus). Animal Behaviour, 31(4), 1237–1243.

Hawkins, G. L., Hill, G. E., & Mercadante, A. (2012). Delayed plumage maturation and delayed reproductive investment in birds. Biological Reviews, 87(2), 257–274.

Hurlbert, S. H. (1984). Pseudoreplication and the Design of Ecological Field Experiments. Ecological Monographs, 54(2), 187–211.

Kappes, P. J., Dugger, K. M., Lescroël, A., Ainley, D. G., Ballard, G., Barton, K. J., Lyver, P. O., & Wilson, P. R. (2021). Age-related reproductive performance of the Adélie penguin, a long-lived seabird exhibiting similar outcomes regardless of individual life-history strategy. Journal of Animal Ecology, 90(4), 931–942.

Kentie, R., Shamoun-Baranes, J., Spaans, A. L., & Camphuysen, K. (2023). Spatial patterns in age-and colony-specific survival in a long-lived seabird across 14 contrasting colonies. Ibis, 165(1), 82–95.

Kralick, A. E., O’Connell, C. A., Bastian, M. L., Hoke, M. K., Zemel, B. S., Schurr, T. G., & Tocheri, M. W. (In review). Body Size Variation among Adult Males and its Implications for Sexual Dimorphism in Orangutans (Pongo spp.). Integrative and Comparative Biology.

Lack, D. (1954). The Natural Regulation of Animal Numbers. Clarendon Press.

Lack, D. (1968). Ecological adaptations for breeding in birds. Methuen.

Lowe, J. R., Russ, G. R., Bucol, A. A., Abesamis, R. A., & Choat, J. H. (2021). Geographic variability in the gonadal development and sexual ontogeny of HEMIGYMNUS, CHEILINUS and OXYCHEILINUS wrasses among INDOLJPACIFIC coral reefs. Journal of Fish Biology, 99(4), 1348–1363.

MacLean, A. A. E. (1986). Age-Specific Foraging Ability and the Evolution of Deferred Breeding in Three Species of Gulls. The Wilson Bulletin, 98(2), 267–279. JSTOR.

McDonald, D. B. (1993). Delayed Plumage Maturation and Orderly Queues for Status: A Manakin Mannequin Experiment. Ethology, 94(1), 31–45.

Nisbet, I. C. T., Weseloh, D. V., Hebert, C. E., Mallory, M. L., Poole, A. F., Ellis, J. C., Pyle, P., & Patten, M. A. (2017). Herring Gull (Larus argentatus). In P. G. Rodewald (Ed.), Birds of North America. Cornell Lab of Ornithology.

Parsons, J. (1971). Cannibalism in herring gulls. British Birds, 64(12), 528–537.

Paynter, R. A. (1947). The fate of banded Kent Island herring gulls. Bird-Banding, 18(4), 156– 170.

Paynter, R. A. (1966). A new attempt to construct life tables for Kent Island Herring Gulls. Bulletin of the Museum of Comparative Zoology at Harvard College, 133, 489–528.

Pickering, S. P. C. (1989). Attendance patterns and behaviour in relation to experience and pair- bond formation in the Wandering Albatross Diomedea exulans at South Georgia. Ibis, 131(2), 183–195.

Pollet, I. L., Bond, A. L., Hedd, A., Huntington, C. E., Butler, R. G., & Mauck, R. (2021). Leach’s Storm-Petrel (Hydrobates leucorhous). In Birds of the World (Version 1.1.). Cornell Lab of Ornithology.

Pyle, P. (2008). Identification guide to North American birds. Part II. Anatidae to Alcidae. Slate Creek Press.

Pyle, P., Nur, N., Sydeman, W. J., & Emslie, S. D. (1997). Cost of reproduction and the evolution of deferred breeding in the western gull. Behavioral Ecology, 8(2), 140–147.

R Core Team. (2021). R: A language and environment for statistical computing. R Foundation for Statistical Computing.

Riotte-Lambert, L., & Weimerskirch, H. (2013). Do naive juvenile seabirds forage differently from adults? Proceedings of the Royal Society B: Biological Sciences, 280(1768), 20131434.

Schaedler, L. M., Taylor, L. U., Prum, R. O., & Anciães, M. (2021). Constraint and Function in the Predefinitive Plumages of Manakins (Aves: Pipridae). Integrative and Comparative Biology, 61(4), 1363–1377.

Shlepr, K., Ronconi, R., Hayden, B., Allard, K., & Diamond, A. (2021). Estimating the relative use of anthropogenic resources by Herring Gull (Larus argentatus) in the Bay of Fundy, Canada. Avian Conservation and Ecology, 16(1).

Taborsky, M., Oliveira, R. F., & Brockmann, H. J. (2008). The evolution of alternative reproductive tactics: Concepts and questions. In R. F. Oliveira, M. Taborsky, & H. J. Brockmann (Eds.), Alternative Reproductive Tactics (1st ed., pp. 1–22). Cambridge University Press.

Tickell, W. L. N. (1968). The Biology of the Great Albatrosses, Diomedea exulans and Diomedea epomophora. In O. L. Austin, Jr. (Ed.), Antarctic Bird Studies (Vol. 12, pp. 1–56). American Geophysical Union.

Tinbergen, N. (1953). The herring gull’s world: A study of the social behaviour of birds. Collins.

Welklin, J. F., Lantz, S. M., Khalil, S., Moody, N. M., Karubian, J., & Webster, M. S. (2021). Social and abiotic factors differentially affect plumage ornamentation of young and old males in an Australian songbird. Animal Behaviour, 182, 173–188.

Wickham, H., Averick, M., Bryan, J., Chang, W., McGowan, L., François, R., Grolemund, G., Hayes, A., Henry, L., & Hester, J. (2019). Welcome to the Tidyverse. Journal of Open Source Software, 4(43), 1686.

Wolfe, J., Ryder, T., & Pyle, P. (2010). Using molt cycles to categorize the age of tropical birds: An integrative new system. Journal of Field Ornithology, 81, 186–194.

Wooller, R. D., & Coulson, J. C. (1977). Factors Affecting the Age of First Breeding of the Kittiwake Rissa tridactyla. Ibis, 119(3), 339–349.

Zeileis, A., Wiel, M. A., Hornik, K., & Hothorn, T. (2008). Implementing a class of permutation tests: The coin package. Journal of Statistical Software, 28(8), 1–23.

