## Supplementary Material for "Adolescent gulls have the opportunity for social development at breeding colonies"

Liam U. Taylor

**SUPPLEMENTARY MATERIAL**

***Table of contents***

pp.	2-3	Count error correction

pp.	4-6	Qualitative observations of social engagement

p.	7	Observations of a predefinitive (S3) parent

**COUNT ERROR CORRECTION**

Counting gulls while walking can be prone to observation error, either because of flushing birds, miscounting large groups, or general chaos. Systematic miscounting that differed between plumage classes could impact my conclusions (e.g., not only do birds flush away before I count them, but also S3 birds flush more readily than Definitive birds). I assessed systematic miscounting by recounting East Beach in the opposite direction (northbound, “reverse”) immediately following a standard (southbound, “forward”) count. Of 68 pairs of counts, 24 were full counts that differentiated S1&2, S3, and Definitive birds in both forward and reverse directions. The remaining 44 pairs were full counts in the forward direction, but “split” counts in the reverse direction of either only S1-3 birds (20 reverse counts) or total birds (24 reverse counts). These split reverse counts were designed to assess territorial proportions (see Main Text *Methods: Social engagement*). I thus had 44 recount pairs for S1-3 birds (24 full reverse, 20 split reverse), 24 recount pairs for Definitive birds (24 full reverse), and 24 recount pairs for total birds (24 split reverse). I assessed systematic miscounting with paired T-tests comparing forward and reverse counts of S1&2, S3, Definitive, and total birds.

There were no significant differences between paired, forward *vs.* reverse counts for S1&2 or Definitive birds (Fig. A1). In contrast, there was a significant difference when comparing forward *vs.* reverse S3 counts (Fig. A1). I conclude that S3 birds may have been slighlty more likely to flush off the counting area during standard (forward) counts, or perhaps repeated inspection helped uncover additional S3 birds.

There was a larger significant difference (~50 individuals) when comparing forward *vs.* reverse Total counts (Fig. A1). This comparison hints at the consequence of counting method. For the Total comparison, forward counts were “full” counts involving plumage identification, whereas reverse counts were “split” counts of Total birds near the water or the berm (see Main Text *Methods: Social engagement*). S1&2 and S3 comparisons also included split reverse counts, but those counts required plumage identification in both directions. In other words, reverse Total counts were the only kind that did not involve searching for predefinitive birds. The significant difference in forward *vs.* reverse Total counts thus suggests I missed birds while focusing on plumage. Exact quantification is impossible, but I suspect I was more likely to gloss over Definitive birds than S1-3 birds.

In conclusion, I found evidence of minor undercounting for S3 individuals and separate evidence of effort-based Total undercounting. These two sources of bias are both small and in opposing directions (<3 S3 individuals undercounted, ~50 Total individuals undercounted, the latter vaguely biased towards Definitive birds). Lacking further details, I make no adjustements to my final estimates.

Two outside observers helped confirm my counts were generally accurate. On July 5, 2022, I walked East Beach with an undergraduate researcher (M. Stretch). I counted 525 total birds in the forward direction (starting 1:07 PM) and 588 in the reverse direction (1:28 PM), while she counted 425 (-100) forward and 479 (-109) reverse. The next day, I repeated this process with a graduate researcher (H. Spina). My counts were 565 forward (4:19 PM) and 538 reverse (5:08 PM) while hers were 531 (-34) forward and 508 (-30) reverse. Comparing these values suggests: (A) experienced observers count more birds (B) some of the difference between forward and reverse counts reflects actual bird movements.

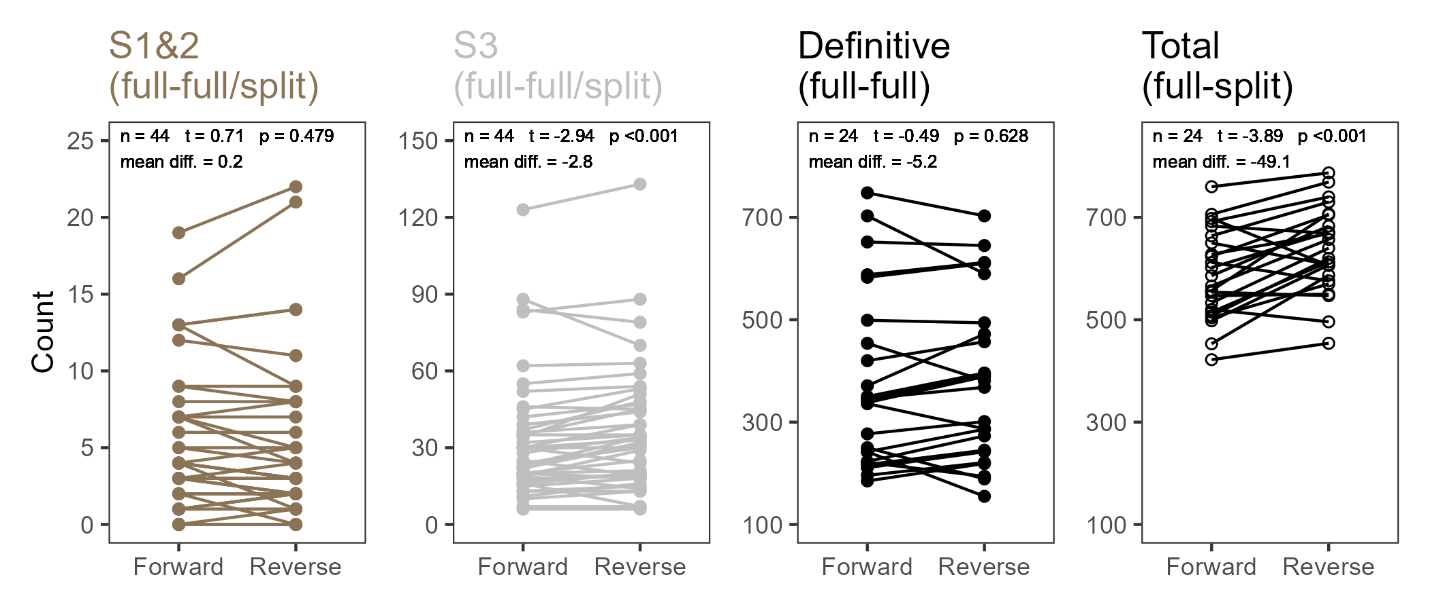
**Figure A1.** Paired counts to assess systematic errors in plumage class estimates of American Herring Gulls on East Beach, Kent Island. Each panel shows pairs of forward and reverse counts. Inset values give results from paired T-tests. Forward counts were all “full” counts, which gave separate values for S1-3 and Definitive birds. Some reverse counts were “split” counts, whereby only S1-3 or Total birds were counted. Split counts were designed to differentiate between birds on territorial *vs.* non-territorial areas (see Main Text *Methods: Social engagement*), but values from both areas were summed for this analysis.

**QUALITATIVE OBSERVATIONS OF SOCIAL ENGAGEMENT**

*Territoriality*

Focal and incidental observations of predefinitive American Herring Gulls (*Larus argentatus smithsonianus*) emphasized the ways in which these birds were engaged in the social context of the breeding colony despite neither holding territories nor breeding. I frequently saw predefinitive Stage 3 (S3) birds (see Main Text *Methods: Plumage stage classification*) standing in territorial areas, sometimes <1 m from active nests. Left undisturbed, some S3 birds would remain loafing, preening, and sleeping in a foreign territory for upwards of 30 min. Others were driven off in seconds or prevented from landing altogether. Activity on foreign territories ended in one of two ways: (1) the S3 bird simply flying away, often far over the island or water, or (2) a Definitive bird charging at the S3 bird, the latter jumping to an adjacent territory before being quickly driven off by a different Definitive bird. It was thus common to see S3 birds ping-ponging among territories. I saw no evidence of a loafing “club” for either predefinitive or definitive birds on either Kent Island or Great Duck Island (Tinbergen, 1953).

On multiple occasions, S3 birds showed they were capable of driving Definitive or other S3 birds out of position. Like most conflicts, these displacements were done with a slow head-down charge or, less frequently, an open-winged lunge (note I mean “displacement” here as “pushing out of place,” not to be confused with Tinbergen’s theory of internal behavioral relations; Tinbergen, 1953). However, the S3 birds that displaced definitive birds from territorial areas were invariably displaced themselves. For example, on July 20, 2022, Kent Island, I saw an S3 bird (Bird P1) drive off a definitive bird (Bird D1) with an open-winged lunge amidst a pack of nests. Just one minute later, a different definitive individual (Bird D2) approached with its head down and bill open, pushing Bird P1 back several steps. Over the next two minutes, Bird D2 slowly drove Bird P1 towards the territory of yet another definitive bird (Bird D3) whose charge was sidestepped by Bird P1. After a minute of no further interaction, Bird P1 softly flew off towards the intertidal.

Although physical contact was less common than symbolic aggression (Tinbergen, 1953), it was clear that social engagement at the breeding colony carried physical risks. For example, on July 17, 2022, Great Duck Island, I witnessed an S3 American Herring Gull stuck in a rocky crevice and beaten into submission by two definitive Great Black-backed Gulls (*L. marinus*) near their nesting territory. Struggling free after 30 s, the S3 bird collapsed on a nearby rock where its flailing drew attacks from other nesting American Herring Gulls. The young bird eventually regained its balance and stumbled towards the water with a clearly broken wing.

Except for the one S3 parent on Kent Island (see below), I saw no good evidence that S3 birds held territories. On the morning of June 15, 2022, Kent Island, one S3 bird was hesitant to flush from a large rock close to the wrack line, instead speed walking ahead of me and giving nervous *kek-kek* alarm calls familiar from nesty birds (Nisbet et al., 2017; Tinbergen, 1953). The same bird gave the same response as I approached later in the afternoon and on the next morning. I never saw the bird again after June 16, nor did I witness any clutch near the rock. Towards the end of June 2022, Kent Island, I began seeing S3 birds standing and flying with seaweed or grass in their bills. This observation suggests that at least some young birds were either (A) partially engaged in nest-building behaviors late in the breeding season or (B) increasingly coming away from conflicts that featured territorial grass-pulling behaviors (Tinbergen, 1953).

*Vocal behavior*

I confirmed that predefinitive birds have a repertoire of social vocalizations (Nisbet et al., 2017). I saw S3 birds giving alarm-related *kek-kek* and *keow* calls, conflict- or announcement-related trumpet calls, conflict- or display-related choking calls, and active copulation calls. I never witnessed a predefinitive bird performing the courtship-related head-toss call or the *mew* call associated with courtship and pair-bonding. I also saw an S2 bird giving a trumpet call while flying above the lighthouse on Great Duck Island. This S2 call was notably higher than the trumpet calls of both S3 and Definitive birds. On the other hand, on August 10, 2022, Kent Island, I saw a two-month-old yearling give *kek-kek* and *keow* calls in a low, adult voice.

Vocalizations were socially relevant for young birds. On July 18, 2022, Great Duck Island, an S3 Herring Gull (Bird P1) was sleeping with its head tucked 1 m from a preening Definitive Herring Gull (Bird D1) on a shed roof. After five minutes, another S3 bird (Bird P2) landed 8 m away from the duo on a different section of the roof. Bird D1 became alert, trumpeting 30 s later. At this trumpet call, Bird P1 became alert and began preening. Two minutes later, Bird P1 gave a full trumpet call, causing another Definitive gull (Bird D2) from across the roof to give a slow head-down charge that displaced Bird P1 by 1 m. In sum, predefinitive birds both respond to trumpet calls (becoming alert, trumpeting in return) and elicit responses via trumpet calls (drawing attention and conflict from older birds).

*Courtship*

Early in the summer, I witnessed moments of sloppy or incomplete courtship behavior between predefinitive and Definitive birds. On May 31, 2022, Kent Island, an S3 bird (Bird P1) stood within 1 m of a Definitive pair (Birds D1 and D2) in a territorial area. Bird D1 performed a series of short head-toss calls while bowing and approaching Bird P1. Bird P1 moved away in response. Bird D2 began performing a sustained bowing and bill-pecking display towards Bird D1. Bird D1 gave a trumpet call, circling Bird D2, at which point Bird P1 turned to approach the pair. Bird D2 then charged Bird P1, the latter flying out to sea while D1 and D2 sat loafing as a pair. Four days later, I saw a pair of Definitive birds (presumably D1 and D2) loafing and courting together on the same rock.

On June 1, 2022, Kent Island, I heard harsh, repeated copulation calls and turned to see an S3 bird (Bird P1) mounting a Definitive bird (Bird D1). Bird P1 was calling, flapping, and tail-wagging as during full copulation. Bird D1 had only a partially raised tail and appeared perturbed while nipping at Bird P1. After a few seconds of this partial copulation, another Definitive bird (Bird D2) made a flying charge from several meters up the beach to drive Bird P1 off the back of Bird D1. Bird D2 then landed and gave a trumpet call. Yet another S3 bird (Bird P2) pushed Bird P1 even further away with a walking charge. Birds P1 and D2 then departed, with Bird D1 approaching Bird P2 with a gentle bill touch and head-toss. Bird P2 returned one head-toss but otherwise did not respond. Bird P2 flew off 30 s later, with Bird D1 departing 30 s after that.

*Predation*

Tinbergen (1953) noted that some young Herring Gulls may specialize in eating eggs from foreign nests. Only once, early in the season, did I see a predefinitive bird actively disrupt a nest. On May 31, 2022, on Kent Island, I flushed several birds off their nests while walking down a narrow section of beach. During the commotion, an S3 bird (Bird P1) stumbled out of the thick beach grass and awkwardly nabbed one egg from an unattended nest. The putative Definitive parent of the nest (Bird D1) then jumped over to Bird P1, causing Bird P1 to drop the egg on a rock. Birds P1 and D1 stood 2 m apart in silence, smashed egg between them. After 5 min, Bird D1 charged Bird P1, the latter retreating 1 m. Bird D1 stood pecking at the smashed eggshell.

**References**

Tinbergen, N. (1953). The herring gull’s world: A study of the social behaviour of birds. Collins.

**OBSERVATIONS OF A PREDEFINITIVE (S3) PARENT**

There was only one American Herring Gull nest on Kent Island with an active S3 parent. This nest (dubbed “β nest”) was on the woody edge of gull nesting habitat, ~5 m from a trickling freshwater stream in the South Basin area of Kent Island (see Main Text Fig. 2). Two other active nests sat within ~10 m. Groups of 1-40 gulls and families of Canada Geese (*Branta canadensis*) would often land at the stream to drink. Parents of the three nests bordering the stream, including the S3 parent of β nest, would frequently charge at gulls wandering too close to territory boundaries. I first noticed the parents of β nest on June 2, 2022. The S3 parent of β nest was larger than the Definitive parent (thus putatively male; Robertson et al., 2016; Tinbergen, 1953). On July 10, I found the actual nest location of β nest, which contained three eggs. On June 30, one of these eggs disappeared. Both parents were actively attending the nest through my departure from Kent Island on July 7.

I noted β nest attendance on 30 occasions between June 2 and July 7: both parents were in attendance nine times (30%), the S3 parent was attending alone nine times (30%), and the Definitive parent was attending alone 12 times (40%). I observed each parent incubating on occasions when the other parent was also attending the nest (e.g., calling from a nearby tree).

When I next visited South Basin on July 22, β nest was decomposed with no parents or chicks in sight. After seeing neither parent for three days, I saw the S3 parent of β nest a few meters behind the former nest site on July 29. On July 31, I saw a Definitive bird nervously attending the same site and noted movement consistent with a single chick in a bush below. I never saw either β parent or any of their chicks again, despite 11 subsequent checks August 1-14. In sum, the S3 (perhaps male) and Definitive (perhaps female) parents of β nest incubated three eggs, lost one egg before hatch, and probably attended at least one chick for at least a week.
